# Normal CA1 place fields but discoordinated network discharge in a Fmr1-null mouse model of fragile X syndrome

**DOI:** 10.1101/152462

**Authors:** Fraser Todd Sparks, Zoe Nicole Talbot, Dino Dvorak, Bridget Mary Curran, Juan Marcos Alarcon, André Antonio Fenton

## Abstract

Silence of FMR1 causes loss of fragile X mental retardation protein (FMRP) and dysregulated translation at synapses, resulting in the intellectual disability and autistic symptoms of Fragile X Syndrome (FXS). Synaptic dysfunction hypotheses for how intellectual disabilities like cognitive inflexibility arise in FXS, predict impaired neural coding in the absence of FMRP. We tested the prediction by comparing hippocampus place cells in wild-type and FXS-model mice. Experience-driven CA1 synaptic function and synaptic plasticity changes are excessive in Fmr1-null mice, but CA1 place fields are normal. However, Fmr1-null discharge relationships to local field potential oscillations are abnormally weak, stereotyped, and homogeneous; also discharge coordination within Fmr1-null place cell networks is weaker and less reliable than wild-type. Rather than disruption of single-cell neural codes, these findings point to invariant tuning of single-cell responses and inadequate discharge coordination within neural ensembles as a pathophysiological basis of cognitive inflexibility in FXS.

## Introduction

Fragile X syndrome (FXS) is a neurodevelopmental disorder, and is the most common inherited form of intellectual disability and single-gene cause of autism spectrum disorders (ASD) characterized by inflexible behaviors. FXS results from silencing the X-linked FMR1 gene and subsequent loss of its protein product fragile X mental retardation protein (FMRP) (Colak et al., 2014; Pieretti et al., 1991). FMRP is a negative regulator of protein synthesis, and because it binds to over 400 mRNAs, the consequences of FMRP loss are diverse and not fully characterized (Brown et al., 2001; Darnell et al., 2011).

Synaptic dysfunction is the dominant hypothesis to explain the intellectual disability associated with FMRP absence, but these studies of synaptic function have only been performed in brain slices from task-naïve animals (Bear et al., 2004; Waung and Huber, 2009). FMRP absence in Fmr1-null mice dysregulates translation of mRNAs that encode dendrite-localized proteins that contribute to synaptic development and function in both pre- and post-synaptic sites; resulting in abnormal dendritic morphology likely due to altered levels of scaffold proteins and glutamate receptors in postsynaptic densities (Bassell and Warren, 2008; Bhakar et al., 2012; Braun and Segal, 2000; Comery et al., 1997; Deng et al., 2011; Patel et al., 2013). One of the best-characterized forms of FMRP-related synaptic dysfunction in brain slices from task-naïve Fmr1-null mice, is excessive group 1 mGluR-stimulated long-term depression (mGluR-LTD), indicating that some abnormalities associated with FMRP loss depend on neural activity (Bear et al., 2004; Dolen et al., 2010; Huber et al., 2002). Although alterations in hippocampal long-term potentiation (LTP) have been elusive (Godfraind et al., 1996), abnormal features of LTP in hippocampus and neocortical circuits have also been reported in brain slices from task-naïve mice after reducing or eliminating FMRP (Chen et al., 2014; Hu et al., 2008; Hunsaker et al., 2012; Larson et al., 2005; Lauterborn et al., 2007; Yun and Trommer, 2011). Abnormal activity-dependent synaptic physiology in task-naïve Fmr1-null mice, as well as other mutant models of syndromic forms of ASD, have led synaptic dysfunction hypotheses to predict disrupted tuning of the discharge of individual neurons to represent information that underlies intellectual ability (Zoghbi and Bear, 2012), which we will here call the “disruption hypothesis.” However, learning and memory are relatively normal in Fmr1-null animals (Bakker et al., 1994; Bhattacharya et al., 2012; Brennan et al., 2006; D’Hooge et al., 1997; Kooy et al., 1996; Zhao et al., 2005), highlighting the gap in understanding how synaptic dysfunction, measured in task-naïve mice, is related to intellectual disability in FXS and similar disorders. Accordingly, we take a systems approach to understand how FMRP loss causes intellectual disability by studying the “missing middle” level of biological function (Laughlin et al., 2000), assessed by the electrophysiology of synaptic populations (local field potentials, LFPs) and the action potential discharge in ensembles of single neurons.

Like FXS patients (Hooper et al., 2008), Fmr1-null mice exhibit cognitive inflexibility (Chen et al., 2014; Kooy et al., 1996; Krueger et al., 2011). This inflexibility has been measured using dorsal hippocampus-sensitive and synaptic plasticity-dependent active place avoidance task variants that test the ability to discriminate between established and novel spatial memories (Cimadevilla et al., 2001; Hsieh et al., 2017; Kubik and Fenton, 2005; Pastalkova et al., 2006; Pavlowsky et al., 2017; Tsokas et al., 2016). Fmr1-null and wild-type (WT) mice learn and remember equally well, the stationary location of a mild foot-shock on a slowly rotating arena in the standard active place avoidance task variant. However, the Fmr1-null mice are impaired when required to discriminate between long-term memory of the initially-learned shock location and the current shock location after the shock zone is either changed or eliminated (Radwan et al., 2016). At the same time, the spectral content of concurrently-recorded LFPs along the dorsal hippocampal somatodendritic axis is indistinguishable between Fmr1-null and WT mice, but the somatodendritic organization of phase-amplitude coupled (PAC) 30- 100 Hz gamma oscillation amplitudes and ~8 Hz theta oscillation phases is abnormal. The aberrations are extreme when cognitive inflexibility manifests in memory conflict and memory extinction tests of flexibility; whereas WT PAC is attenuated when the shock zone is changed or eliminated, Fmr1-null PAC persists (Radwan et al., 2016). These observations suggest an alternative “hyperstable/discoordination” hypothesis, that neural information processing is excessively stable and consequently poorly coordinated in the absence of FMRP, leading to inflexible and poorly coordinated neural representations and behavior. We evaluated the disruption and the hyperstable/discoordination hypotheses, which make different predictions for how loss of FMRP affects neural information processing.

To evaluate predictions of the neural information disruption and hyperstability/discoordination hypotheses, we used the neural representation of location in the discharge of hippocampus place cells as a model of flexible cognitive information processing (Fenton, 2015a; Kelemen and Fenton, 2016; Moser et al., 2015; O’Keefe and Nadel, 1978). Whereas the disruption hypothesis asserts altered Fmr1-null hippocampus place coding such as poor quality place fields, the hyperstable/discoordination hypothesis predicts normal place coding that is abnormally indifferent to environmental conditions.

## Results

### Exaggerated baseline and learning-induced changes of synaptic function in Fmr1-null mice

We began by examining synaptic function in the gene knockout Fmr1-null mouse using *ex vivo* hippocampal slice physiology. In agreement with prior work (Franklin et al., 2014; Godfraind et al., 1996; Hu et al., 2008; Lauterborn et al., 2007), we find that CA3 Schaffer collateral to CA1 synaptic efficacy and potentiation does not differ between WT and Fmr1-KO brain slices taken from task-naïve mice (Fig. 1). We then tested whether synaptic function of task-experienced mice differs, measured one day after memory and control training. Fmr1-KO mice performed as well as WT mice (Fig. S1) in the hippocampus- and LTP-dependent active place avoidance task (Cimadevilla et al., 2001; Pastalkova et al., 2006), replicating a prior report (Radwan et al., 2016). We observed training-induced changes in synaptic function, consistent with prior findings using extended training protocols (Park et al., 2015; Pavlowsky et al., 2017). Specifically, greater synaptic efficacy was observed in the trained WT group compared to the home cage group as well as the exposed WT control group that experienced the training environment but were never shocked (Fig. 1A). Synaptic responses from the exposed Fmr1-KO group were almost twice as large as the task-naive Fmr1-KO and the WT groups (Fig. 1B); synaptic responses in the Fmr1-KO trained group were also enhanced, similar to the exposed Fmr1-KO group (Fig. 1B). Synaptic potentiation after 100-Hz high frequency stimulation was indistinguishable between the WT and Fmr1-KO task-naïve home cage groups, as previously reported (Godfraind et al., 1996; Hu et al., 2008). Potentiation was also similar in the WT task-naïve and exposed control groups and potentiation in these groups was greater than the potentiation in the WT trained group (Fig. 1C), as has been reported after extended training (Pavlowsky et al., 2017). The early and late phases of the potentiation were increased in the exposed Fmr1 KO group compared to the mutant task-naive and trained groups, as well as the WT groups (Fig. 1C,D). Moreover, the difference in the amplitude of synaptic potentiation between the WT trained and exposed groups (Fig. 1C) was substantially smaller than the difference between the Fmr1 KO trained and the exposed and task-naive mutant mice (Fig. 1D). These observations indicate that experience-dependent CA1 synaptic function changes are enhanced in Fmr1 KO animals and that experience-driven modulation of CA1 synaptic function is intensified in Fmr1-null mice compared to mice that express FMRP.

**Figure 1.**
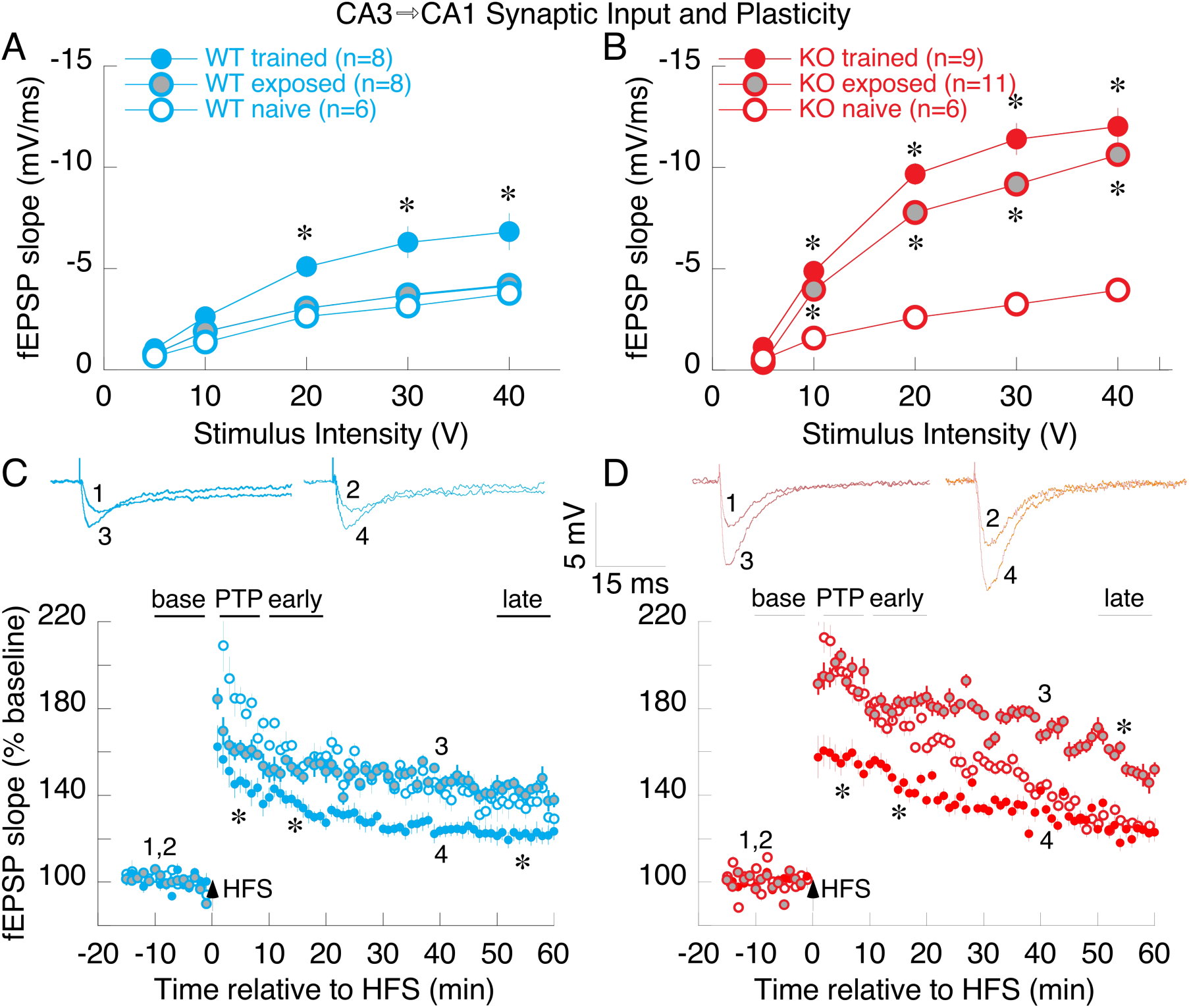
See also Figure S1. Abnormal experience-dependent changes of baseline and plastic hippocampal CA3→CA1 synaptic function in Fmr1-KO mice. A&B) Comparing efficacy of baseline synaptic transmission in WT (A) and Fmr1-KO (B), mice that are either naïve, or after control exposure or memory training in the active place avoidance task. WT and Fmr1 KO synaptic responses are indistinguishable in naïve mice (A,B open circles). Memory training enhances responses in both genotypes (A,B filled colored circles); the enhancement is greater in Fmr1 KO mice, which unlike WT, show enhancement even after control exposure (B, gray circles). Two-way genotype x training ANOVA on the area under the curve confirmed significant effects of training (F_2,42_=25.7, P = 10^-8^) and the genotype x training interaction (F_1,42_=3.49, P = 0.04). Post-hoc Tukey tests confirmed the pattern KO-naïve = WT-naïve = WT-exposed < KO-exposed = KO-trained = WT-trained. C&D) Synaptic potentiation to 100-Hz high-frequency stimulation (HFS) in WT (C) and Fmr1-KO (D) mice. HFS induces post-tetanic potentiation (PTP), early-potentiation, and late-potentiation. Potentiation at each phase appears similar in the naïve WT and Fmr1 KO mice (C,D open circles) and similar in WT naïve and exposed mice (C, open and gray circles, respectively). Potentiation is greater in exposed than naïve Fmr1 KO mice (D, open and gray circles, respectively), but not different between exposed and naïve WT mice (C, open and gray circles respectively). Potentiation is reduced in trained mice of both genotypes (C,D filled colored circles), except late potentiation in trained and task-naive Fmr1 KO mice is not different, but is less that in Fmr1 KO exposed mice. The genotype x training x phase 3-way repeated measures ANOVA on synaptic plasticity showed significant effects of phase (F_3,40_ = 214.2, P = 10^-24^), and the genotype x phase (F_3,40_ = 5.28, P = 0.003) and training x phase (F_680_ = 10.6, P = 10^-8^) interactions so each post-stimulus phase was analyzed separately by 2-way genotype x training ANOVA. The *de novo* protein-synthesis independent PTP and early potentiation changes were greater in exposed Fmr1-KO mice than in exposed WT mice (genotype effects - PTP: F_1,42_ = 7.79, P = 0.008; early potentiation: F_2,42_ = 13.11, P = 0.0008) and smaller in trained compared to naïve and exposed mice for each genotype (training effects - PTP: F_2,42_ = 13.72, P = 10^-5^; early potentiation: F_2,42_= 14.42, P = 10^-5^). *De novo* protein-synthesis dependent late potentiation was not different between the genotypes (F_1,42_= 0.70, P = 0.4) but it was weakest in the trained groups (training effect: F_2,42_ = 19.36, P = 10^-6^). *P < 0.05 relative to task-naive. In C and D, traces 1 and 2 represent baseline fEPSP responses and traces 3 and 4 represent potentiated fEPSP responses from exposed and trained mice, respectively.

### Non-spatial and spatial single-cell discharge features of CA1 place cells are intact in Fmr1-null mice

We then compared the characteristics of dorsal CA1 single unit discharge in freely-behaving WT and Fmr1-KO mice in fixed environments (Fig. S2). Of the 1115 single unit waveform clusters that were identified, 499 were recorded from 12 WT mice and 616 were recorded from 9 Fmr1-KO mice. When the single unit isolation quality was sufficiently high (IsoIBG and IsoINN each > 4 bits) in standard recordings, these were classified as putative principal cells or interneurons. Thus 226 WT and 334 Fmr1 KO presumptive cells comprised the dataset, and were subclassified into functional classes as place cells, non-spatial pyramidal cells, and interneurons. A lower proportion of these were classified as putative interneurons in WT mice (28, 12.4%) than Fmr1-KO mice (98, 29.3%; test of proportions z = 9.4, p = 10^-17^). However, according to standard criteria (Fig. S3) (Fenton et al., 2008; Fox and Ranck, 1975, 1981), when the putative principal cells were classified as place cells (WT n = 108 (47.8%), KO n = 141 (42.2%), the prevalence was lower in the Fmr1-KO mice (test of proportions z = 2.04, p = 0.02). The fundamental extracellular waveform and non-spatial discharge characteristics of the classified cells were indistinguishable between the genotypes (Fig. S3 for firing rate, burst propensity, inter-spike interval, and firing rate comparisons). Furthermore, the fundamental measures of spatial discharge quality did not differ between either the WT and Fmr1 KO place cells or interneurons (Table S1). This finding is unexpected according to disruption hypotheses (Fig. 2).

**Figure 2.**
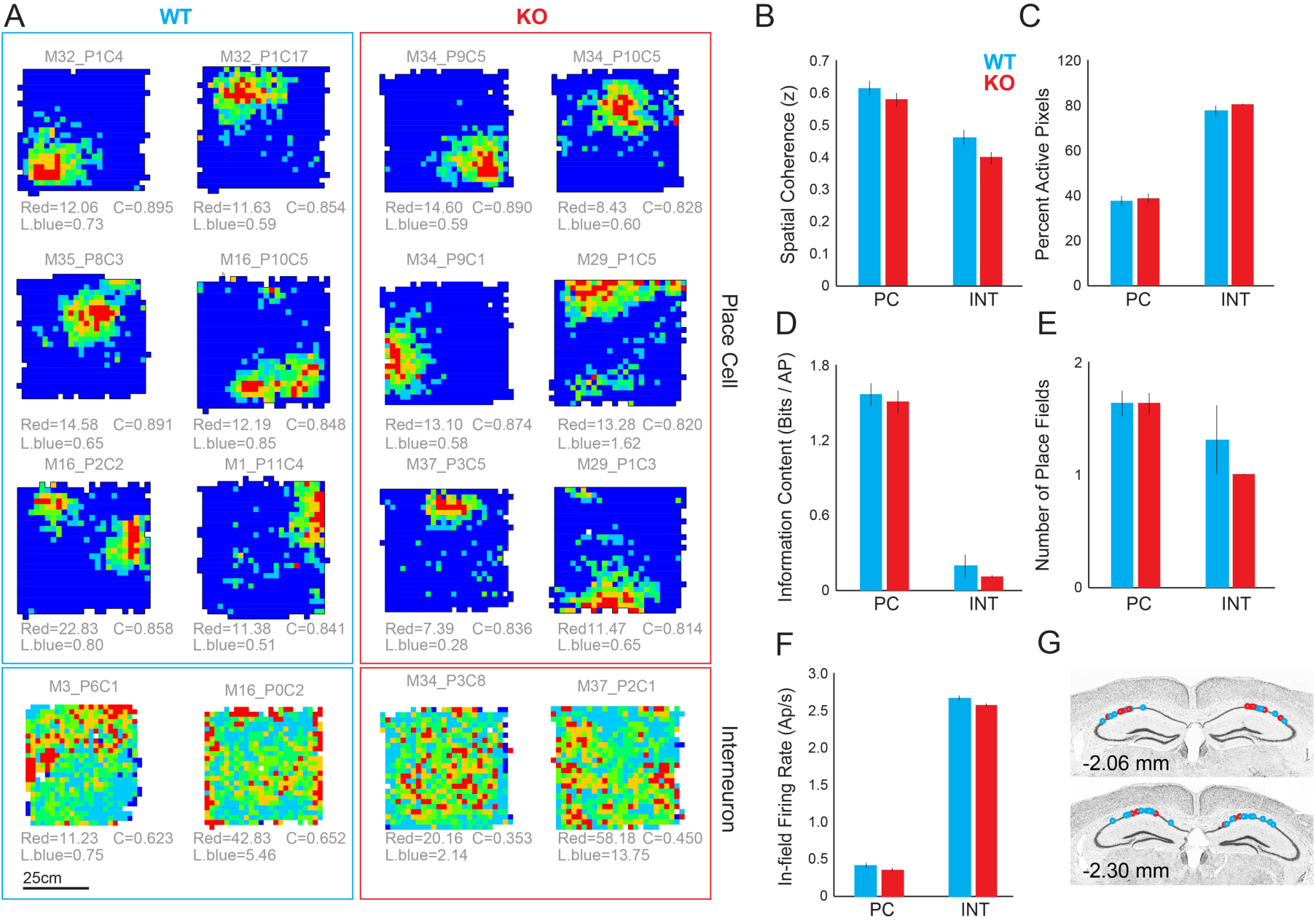
See also Figure S2 and S3. Similar spatial properties of WT and Fmr1-KO CA1 place cells and interneurons in a fixed environment. Place cells and interneurons recorded in the box from the A) WT and B) Fmr1-KO mice. Spatial firing quality differs between place cells and interneurons but not between genotypes as assessed by 2-way genotype x cell class ANOVAs. B) Spatial coherence (genotype: F_1,340_= 0.60; P = 0.44; cell class: F_2,340_= 211.2; P = <0.0001; interaction; F_2,340_= 2.2; P = 0.11). C) Proportion of the environment in which the cell discharged (genotype: F_1,340_= 3.7; P = 0.06; cell class: F_2,340_= 105.2; P = <0.0001; interaction; F_1,340_= 2.3; P = 0.10). D) Information content (genotype: F_1,340_= 3.6; P = 0.06; cell class: F_2,340_= 58.2; P = <0.0001; interaction; F_2,340=_ 2.8; P = 0.06). E) Number of place fields (genotype: F_1,340_= 0.13; P = 0.72; cell class: F_2,340_= 4.0; P = 0.02; interaction; F_2,340_= 2.2; P = 0.11). F) In-field firing rate (genotype: F_1,340_= 0.24; P = 0.62; cell class: F_2,340_=188.6; P < 0.001; interaction: F_2,340_=0.009; P = 0.92). G) WT photomicrograph illustrating the placement of recording electrodes in dorsal CA1.

### Abnormal spike-field coordination in CA1 of Fmr1-null mice

Although the power spectra of LFP oscillations recorded from the CA1 pyramidal cell layer is indistinguishable between the genotypes, the cross-frequency organization of ~8 Hz theta and 30-100 Hz gamma oscillations differs (Radwan et al., 2016), motivating us to examine how oscillations organize the discharge of individual cells (Fig. S4). Oscillations in the LFP organize discharge in cell-specific and frequency-specific patterns (Fig. 3A). While the modulation of discharge in the theta (~8 Hz) and low gamma (30-50 Hz) bands is weaker in Fmr1 KO compared to WT non-spatial pyramidal cells and interneurons, theta modulation but not 30-100 Hz gamma modulation of place cell discharge was similar between the genotypes (Fig. 3B). We then investigated the heterogeneity of the phase-frequency discharge probability relationships amongst cells of each of the three functional classes in the theta and gamma bands by computing the mean-square error (MSE) between the phase-frequency discharge probability histograms of all pairs of cells (Fig. 3C). These relationships were the most stereotyped (corresponding to low MSE) for interneurons in both genotypes while the relationships were the least stereotyped (corresponding to high MSE) for non-spatial pyramidal cells in both genotypes. The relationship was more stereotyped in Fmr1-KO mice compared to WT mice for all cell classes in the theta band (Fig; 3C left; place cells: t_9396=_ 13.44; p = 10^-41^; non-spatial pyramidal cells: t_1031_ = 21.95; p = 10^-88^; interneurons: t_2380_ = 13.79; p = 10^-41^), while in the gamma range, only the non-spatial pyramidal cells and interneurons appeared more stereotyped in Fmr1-KO mice compared to WT mice (Fig. 3C right; place cells: t_9396_ = 1.57; p = 0.12; non-spatial pyramidal cells: t_1031_ = 13.38; p = 10^-37^; interneurons: t_2380_ = 4.34; p = 10^-5^). These data indicate that while individual cells in the Fmr1-null mice are poorly organized by the phase of theta and gamma oscillations in the LFP, the mutant cells are nonetheless stereotyped in the particular ways that spiking is organized by the oscillations that arise from population synaptic activity.

**Figure 3.**
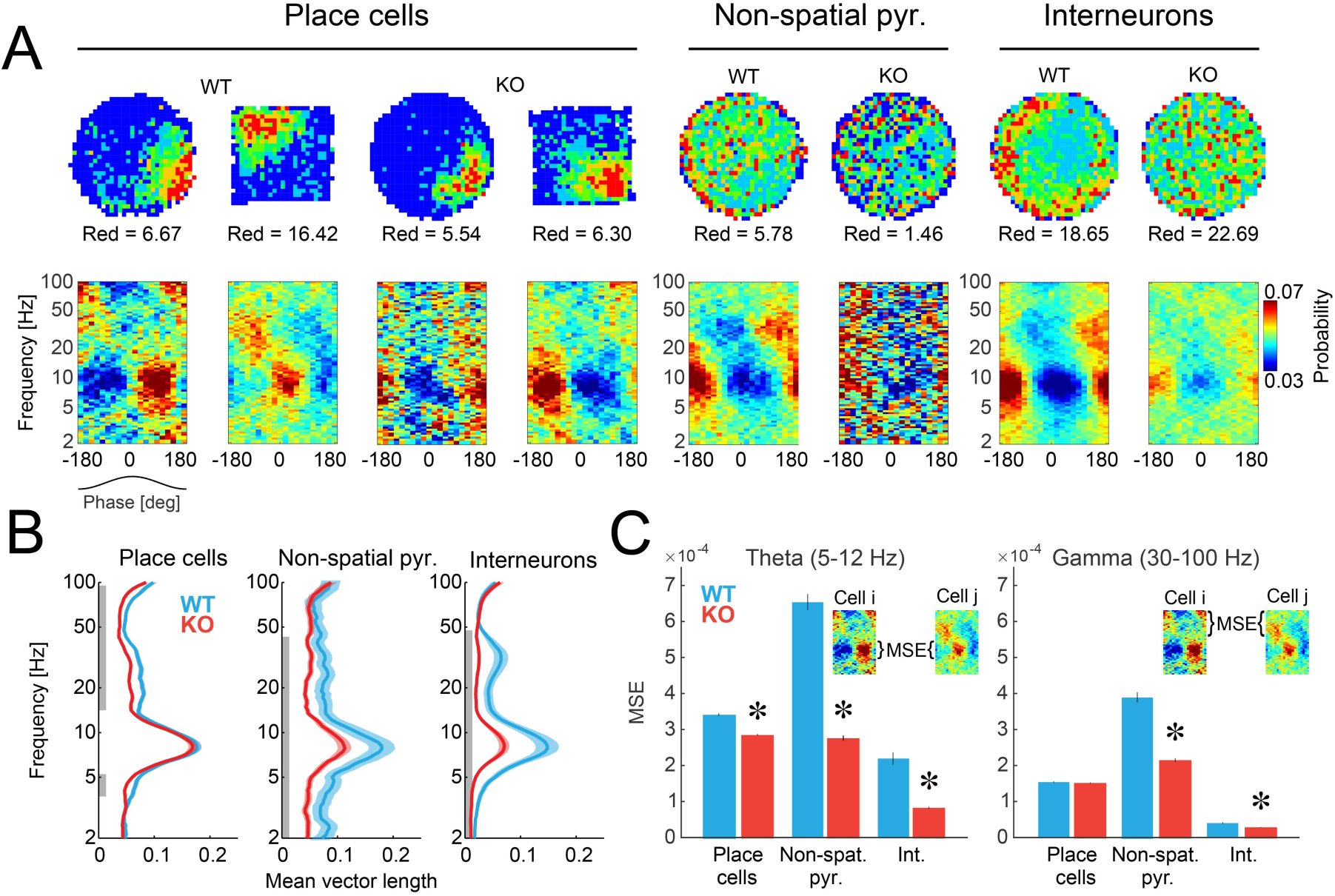
See also Figure S4. Abnormal organization of Fmr1 KO principal cell spiking by LFP oscillations. A) Cell-specific spatial firing rate maps and phase-frequency discharge probability histograms of place cells, non-spatial pyramidal cells, and interneurons. B) Coupling strength between spikes and field measured as mean vector length. The gray line indicates continuous bands with P < 0.05 for at least 3 neighboring frequencies). C) Mean square error (MSE) differences between phase-frequency discharge probability histograms across all pairs of cells for the theta (5-12 Hz; left) and gamma (30-100 Hz; right) frequency bands. Lower MSE indicates reduced variety. *P < 0.05 between genotypes.

### Reduced spatial discharge coordination and weaker network states in the short-time scale discharge dynamics of Fmr1 -null place cells

We next examined the reliability of spatial discharge by measuring overdispersion during passes through the place fields (Fenton et al., 2010; Fenton and Muller, 1998; Jackson and Redish, 2007). The overdispersion of WT mouse place cells in standard environments was comparable to what has been reported for rats (Fenton et al., 2010; Fenton and Muller, 1998; Jackson and Redish, 2007). Overdispersion was lower (spatial firing was more reliable) in Fmr1-KO mice when the unit of analysis is a pass through a place field (WT σ^2^ = 5.81; KO σ^2^ = 4.89; F_6775,9101_ = 1.19, p = 10^-17^) and even when the unit of analysis is a place cell (σ^2^ = 5.98±0.44; KO σ^2^ = 5.02±0.3; t_176_ = 1.87, one-tailed p < 0.03).

This greater spatial discharge reliability of individual Fmr1-KO place cells is accompanied by significantly less spatial discharge covariance of the mutant cell pairs with overlapping place fields (Fig. 4A top right; r = 0.042±0.013) compared to wild type (Figure 4B; r = 0.0917±0.028; t_372_ = 2.12 p<0.03). This difference was due to greater WT discharge covariance in the upper 50% of the distribution (WT r = 0.29±0.03; KO r = 0.21±0.01; t_178_ = 2.94 p = 0.004, whereas the lower halves of the distributions did not differ between the genotypes (WT r = -0.12±0.02, KO r = -0.13±0.01; t_176_ = 0.66 p < 0.51). These findings indicate that loss of FMRP selectively reduces the positively coordinated firing rate fluctuations between place cells on the timescale of the 1-5 seconds that it takes to traverse a firing field (Fenton and Muller, 1998). We therefore examined in a second way, whether Fmr1-KO place cells form weaker network states as indicated by this weak spatial discharge coordination. The location-independent, time scale-specific Kendall’s correlation (τ) between each pair of simultaneously-recorded principal cells directly estimates the strength of the network states (Neymotin et al., 2017; Schneidman et al., 2006). At the 5-s behaviorally-relevant timescale (Fig. 4C), as well as the 25 ms and 40 ms time scales of gamma, Fmr1-KO cell pair discharge correlations were weaker than WT cell pairs (Fig. 4D). The same trend did not reach significance for theta 125 ms (data not shown) and 250 ms time scales. Thus, weaker network states of principal cell discharge result from loss of FMRP. These states are less likely to recur in Fmr1 KO ensembles compared to WT ensembles, estimated by population coordination (PCo; Pearson’s correlation of the vector of Kendall correlations from the two halves of a recording, Fig. 4D) (Neymotin et al., 2017).

**Figure 4.**
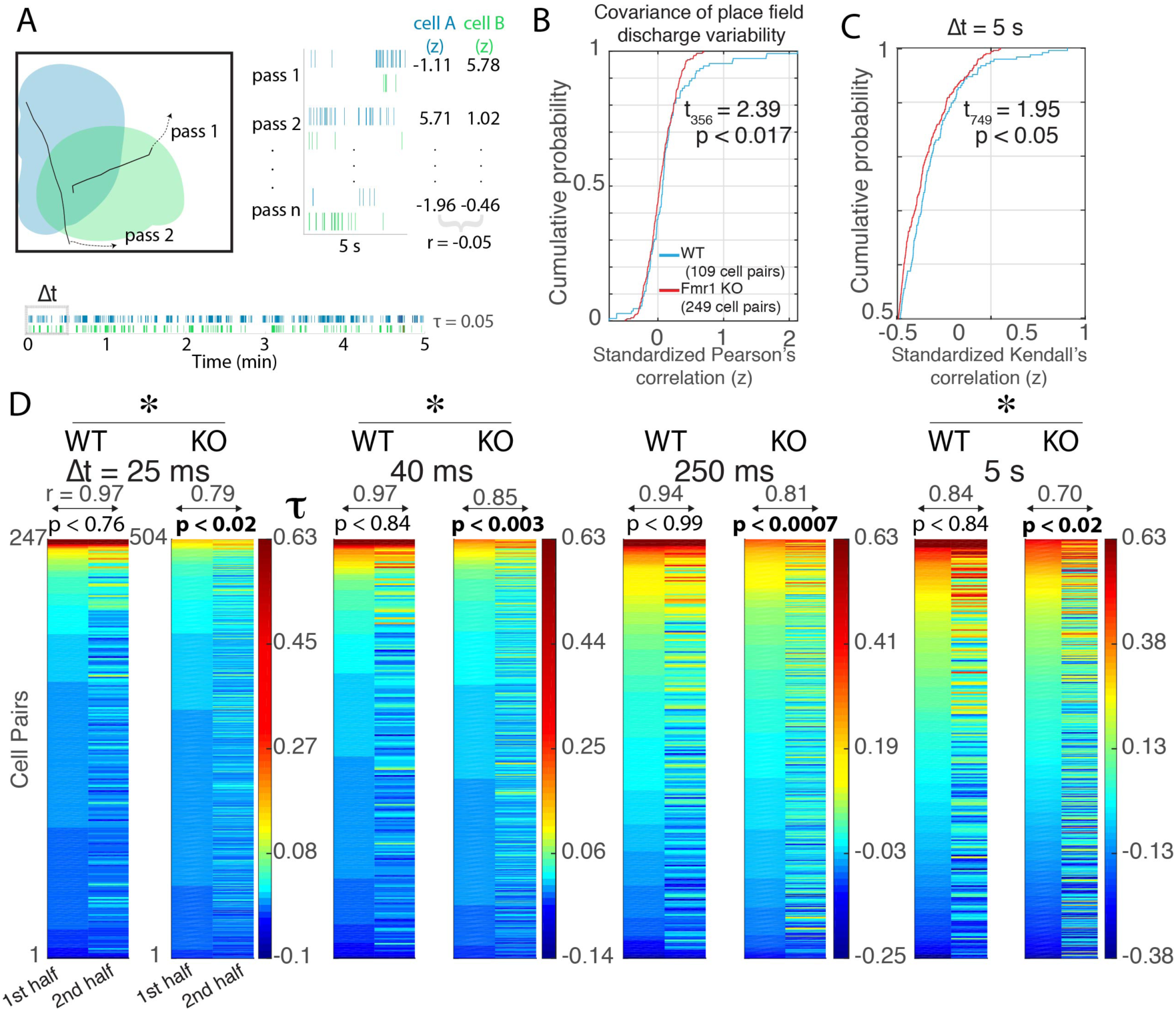
Weak network states of Fmr1-KO place cell ensembles estimated by the set of spike train correlations. A) Schematic illustrating two place cell firing fields that are overlapping, with two 5-s trajectories that pass through both fields. Bottom: 5-min spike rasters from example cells and Right: a 5-s raster subsample with corresponding standardized firing (z) values depicted for n passes. B) Pearson’s correlation of the two sets of z values describes the covariance of the place field discharge variability between the two cells. Fmr1-KO and WT cumulative probability distributions of all the pairwise correlations illustrate reduced covariance of the Fmr1-KO place field discharge variability and C) weaker temporal coupling within place cell ensembles, here assessed at Δt = 5-s time scale. D) Kendall’s correlation (τ) computed at a particular resolution (Δt) estimates coupling of the two spike trains. Each pair of columns depicts the two PCorr vectors of τ between the set of simultaneously recorded place cell spike trains during two 10-min intervals. The correlations from the earlier interval are sorted high to low and the cell-pair identities are preserved across intervals, to illustrate the recurrence of the network state that the vectors estimate. Recurrence is estimated by the PCo, Pearson correlation of the two vectors, which is given at the top of each vector pair. Across the gamma (25 and 40 ms), theta (125 and 250 ms) and behavioral (1 and 5 s) timescales, positive correlations between Fmr1-null place cells tend to be less prevalent and the network state is less likely to recur. The p value for the t test comparison between the two halves of the recording is given at the top of each PCorr pair. The df = 246 and 503 for the WT and KO comparisons, respectively. *P < 0.05 between genotypes

### During dissociation of spatial frames, the fluctuations of spatial-frame specific place cell discharge is more weakly coordinated in Fmr1-null than in wild-type ensembles

Continuous rotation of the circular arena dissociates the environment into the spatial frames of the stationary room and the rotating arena, allowing us to examine the ability of place cell ensembles to dynamically switch between representing locations in the dissociated spatial frames on sub-second timescales (Kelemen and Fenton, 2010, 2016). Although mice in the rotating arena were not performing an explicit navigation or spatial memory task and were not shocked or otherwise manipulated, place cell discharge alternated between signaling locations in the stationary and rotating spatial frames in both genotypes (Fig. 5). The session-averaged frame-specific spatial firing was better organized in the room frame, and this preference was similar for the two genotypes (Fig. S5). Figure 5A illustrates the dynamics of frame-specific discharge during rotation measured as the time series of *ΔI_pos_*. This frame-specificity spontaneously alternates between the stationary and rotating frames and is overall stronger in the Fmr1-KO mice compared to WT (Fig. 5B). Place cells from both genotypes tended to preferentially signal stationary locations over rotating locations.

**Figure 5.**
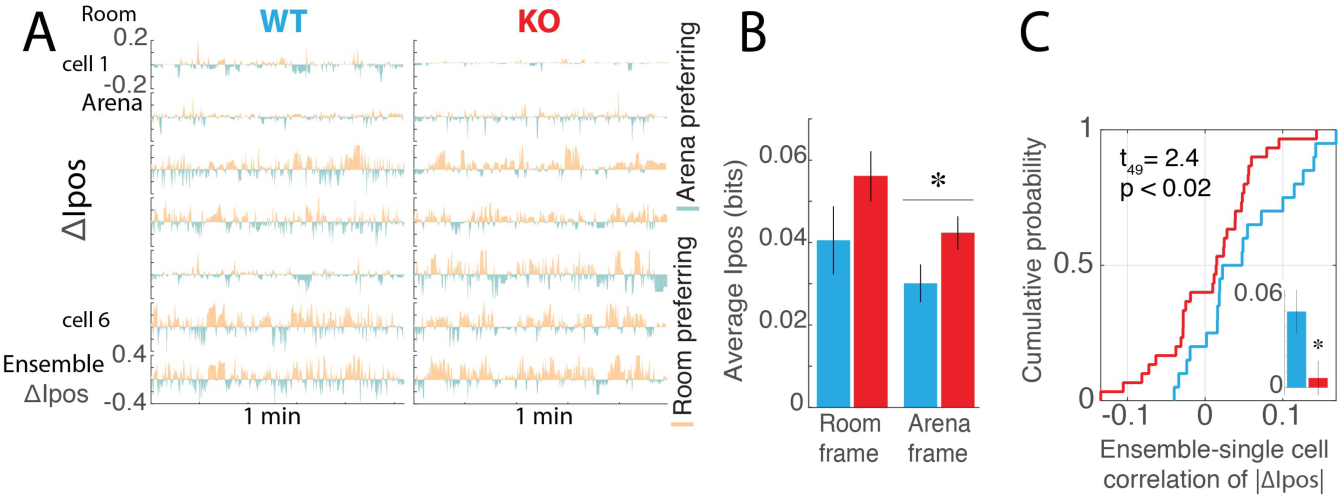
See also Figure S5. Abnormal spatial frame-specific place cell discharge in Fmr1-KO place cells during environmental dissociation. A) WT and Fmr1-KO single-cell examples of *ΔI_pos_* time series illustrating the spatial frame-specific positional information alternation between the stationary room and rotating arena frames. *I_pos_* is computed at 133 ms resolution. Each time series depicts the cell-specific difference between *ΔI_pos_* in the room frame and *I_pos_* in the arena frame (*ΔI_pos_*). The ensemble *ΔI_pos_* time series is depicted on the bottom row. B) Average ± s.e.m. of the room -specific and arena-specific *_Ipos_* values for WT and Fmr1 KO. C) Cumulative probability distributions and average ± s.e.m. (inset) summarizing the correlations of frame-specific positional discharge fluctuations |*ΔI_pos_*| between single place cells and the rest of the ensemble (WT n = 20, KO n = 30). *P < 0.05 between genotypes.

We then examined how well these dynamics were correlated amongst the network of cells by asking how the frame-specific spatial coding dynamics of a single cell is correlated with the frame-specific spatial coding fluctuations of the remaining ensemble of cells. The ensemble fluctuations of frame-specific discharge (|*ΔI_pos_*|) were computed for all simultaneously recorded cells except one, and the correlation of the ensemble and “left-out” single cell |*ΔI_pos_*| time series was calculated. The association between the ensemble and single cells was weaker in Fmr1-KO compared to WT place cells (Fig. 5C), indicating weaker network states in the Fmr1-null mice.

## Discussion

### Experience-dependent synaptic function abnormalities, and intact place cell tuning in Fmr1-null mice

The basic place cell properties that we have measured here are disrupted by a variety of genetic manipulations that affect synaptic plasticity, including deletion of NMDA receptors (Dragoi and Tonegawa, 2013; McHugh et al., 1996; McHugh et al., 2007; Nakazawa et al., 2002; Nakazawa et al., 2003), the kinases PKA (Rotenberg et al., 2000), and CaMKII (Cho et al., 1998; Rotenberg et al., 1996), but effects of Fmr1 KO on hippocampal information processing during behavior have not been previously described. The absence of FMRP dysregulates translation of hundreds of mRNAs, modifying the expression of channel proteins (Brown et al., 2001; Gross et al., 2011), and altering neuronal excitability and synaptic function in hippocampus, and possibly throughout the brain as assessed by *in vitro* studies (Deng et al., 2011; Gibson et al., 2008; Zhang and Alger, 2010; Zhong et al., 2010). Such consequences have motivated disruption hypotheses that predict impaired information processing in the neural networks of individuals without FMRP. However, the present findings appear incompatible with disruption hypotheses and strongly support alternative hyperstable/discoordination hypotheses for the consequences of FMRP loss in a neural circuit leading to intellectual disability in FXS. While we confirmed that Schaffer collateral synaptic transmission and plasticity are indistinguishable between naïve WT and Fmr1 KO brain slices as previously reported (Franklin et al., 2014; Godfraind et al., 1996; Hu et al., 2008; Lauterborn et al., 2007), we also uncovered substantial abnormalities in the mutant mice, but only after spatial experience (Fig. 1). The experience-dependent increase in synaptic transmission and synaptic plasticity were observed 24-h after active behavior in a novel environment as we have shown before in WT mice (Pavlowsky et al., 2017), but the changes were excessive in Fmr1 KO mice; manifest whether or not place learning was conditioned. WT mice only express such changes after conditioning (Fig. 1). This, the first demonstration of these abnormalities of the Fmr1 KO mouse, are consistent with reduced translation repression causing excessive experience-dependent long-term changes in synaptic function. While this has been hypothesized on the basis of exaggerated mGluR-LTD, it has not been generalized to enhanced plasticity (Bear et al., 2004; Zoghbi and Bear, 2012). The present demonstration, one of the first, that synaptic transmission and potentiating synaptic plasticity are abnormally enhanced in the absence of FMRP, highlights the importance and value of incorporating behavioral assessment into studies of brain slices to assess cellular and network parameters to understand mechanisms of behavior and cognition.

This study is also one of the first to extend the investigation of the mechanisms of behavior and cognition in FXS to a freely-behaving mouse model of the disorder. We find that in freely-behaving Fmr1-null mice, the basic single-cell electrophysiological properties of dorsal CA1 neurons are normal. Although we classified more single units as interneurons in Fmr1-null mice and a lower proportion of the putative principal cells as place cells, the place cell discharge properties were nonetheless normal, suggesting complex information processing is also intact (Fig. 2). Indeed, the spatial firing properties of Fmr1-null and WT place cells were indistinguishable, even during the challenge of arena rotation (Fig. S5), dissociating space into two distinct spatial frames. Rotation disturbs rat place cells if there has been no spatial training, as in the present case (Zinyuk et al., 2000). While disruption was not observed in mice (Figs. 5; S5), the similar intact place cell responses of Fmr1-null and WT mice parallels the intact memory and navigation that Fmr1-null mice display (Fig. S1) during place avoidance tasks on the rotating arena (Radwan et al., 2016). Indeed, like others (Bakker et al., 1994; Bhattacharya et al., 2012; Brennan et al., 2006; D’Hooge et al., 1997; Zhao et al., 2005), the most reliable cognitive deficits we have previously detected in Fmr1-null mice is cognitive inflexibility; impaired ability to update learned avoidance of a shock zone when the location of shock changed (Radwan et al., 2016). If one interprets the hippocampus place code as a dedicated code (see (Fenton, 2015a), then these findings directly contradict predictions from disruption hypotheses: place cell cognitive information processing appears normal despite abnormal experience-dependent synaptic function. Importantly, we have argued that the hippocampus place code is not a dedicated code and instead should be interpreted as an ensemble code comprised of neurons with mixed selectivity responses (Fenton et al., 2008; Park et al., 2011; Xie et al., 2016), which emphasizes the temporal interactions in the discharge between cells, called neural coordination and deemphasizes single cell discharge properties like firing rates, place fields in light of multimodal response tuning (Buzsaki, 2010; Fenton, 2015a, b; Fusi et al., 2007; Harris et al., 2003; Phillips and Singer, 1997).

### Neural discoordination in the hippocampus network of Fmr1-null mice

Consistent with alternative hyperstable/discoordination hypotheses that interpret the hippocampus place code as an ensemble code, hippocampal discharge is by no means normal in Fmr1-null mice. The substantial abnormalities argue for a different conceptualization of the pathophysiology that can explain cognitive dysfunction in FXS and autism, and perhaps other conditions that are characterized by cognitive information processing deficits (Phillips and Silverstein, 2003; Uhlhaas and Singer, 2007). The coordinated activity between groups of hippocampal principal cells is abnormal in Fmr1- null mice, indicating a dissociation between intact single cell properties and the temporal coordination of their interactions at the level of the hippocampus information processing network, as predicted by discoordination hypotheses for cognitive dysfunction both generally (Fenton, 2015b; Lee et al., 2014) and in the case of FXS in particular (Radwan et al., 2016). Accordingly, inferences from single cell properties can be misleading, and instead, it may be the temporally-organized interactions amongst cells that provide informative indicators of cognitive function (Buzsaki, 2010; Fenton, 2015a, b; Johnson et al., 2009; Okun et al., 2015; Suh et al., 2013).

Relationships between the discharge of single CA1 cells and oscillations in the LFP generated by synaptic interactions amongst large numbers of cells reveal abnormally weak coupling between individual cells and the population in Fmr1 KO mice (Fig. 3B). Furthermore, the set of these weak relationships is abnormally stereotyped. Within this abnormal network state, we also observed a substantially reduced variety of spike-phase relationships between non-spatially tuned cells and theta oscillations in the Fmr1-null LFP (Fig. 3C). Thus activity within the Fmr1-null hippocampus network of principal cells forms an abnormally invariant and weak spatial firing network infrastructure within which the intact single cell location-specific activity of place cells is embedded. Because principal cells in hippocampus drive inhibition that organizes network dynamics crucial for information processing (reviewed in Buzsaki, 2010), it is likely pathological that in Fmr1-null mice there is an increased prevalence of non-spatial pyramidal cells, with spiking that is poorly organized by the ongoing LFP (Fig. 3B), forming a relatively functionally-homogeneous group of neurons (Fig. 3C). This may be one of the contributors to both the weak network states (Fig. 4) and the paradoxically rigid single place cell properties of Fmr1-null mice (Figs. 2,5).

### System-level origins of intellectual disability in FXS

The interactions between the spike trains of multiple cells provide crucial additional evidence that despite their intact place tuning properties, individual Fmr1-null place cells form abnormally weakly coordinated networks at the time scale of inhibition-associated gamma oscillations, as well as an abnormally weakly coordinated network of cells at multiple longer time scales up to a few seconds (Fig. 5). Together these findings provide a novel conceptual picture of intellectual disability in FXS and the associated autistic features; Fmr1-null neural circuits are comprised of functionally normal individual cells (Fig. 2) that express stereotypically weak discharge relationships to the phases of ongoing oscillations in the LFP and form correspondingly weak network states because they inadequately cohere into common-function neural ensembles (Fig. 4; 5D). This viewpoint can account for findings that basic learning and memory are intact whereas cognitive and memory inflexibility characterize FXS-model mice (Radwan et al., 2016). If correct, this should engender optimism because therapeutic approaches may only need to target the mechanisms of coordination rather than the potentially less easy to manipulate mechanisms that establish and tune the response properties of individual cells.

Going forward, it will be important to identify proximal mechanisms of the discoordination associated with FMRP absence, in the pursuit of novel therapies. Initial clues point to inhibition abnormality in the regulation of spike timing (Anastassiou et al., 2010), more so than increased excitability which may also explain increased seizure likelihood in the absence of FMRP (Dolen et al., 2007; Zhong et al., 2010).

## Experimental Procedures

### Subjects

Wild-type mice with a mixed C57BL6/FVB background were used as well as Fmr1-null mice carrying the Fmr1^tm1Cgr^ allele on the same mixed C57BL6/FVB background. The mutant mice were obtained from Jackson Laboratories (Bar Harbor, ME) to establish local colonies, and housed in a room with a 12-h light/dark cycle (light on at 7 am) with access to food and water *ad libitum*. All experimental procedures were performed as approved by the Institutional Animal Care and Use Committees of NYU and the SUNY, Downstate Medical Center, and according to NIH and institutional guidelines and the Public Health Service Policy on Humane Care and Use of Laboratory Animals.

### Ex vivo slice electrophysiology

Male WT (n = 22) and Fmr1-null mice (n = 26) aged 3-4 months were anesthetized with isoflurane (5% in 100% O_2_ for 3 min) 30 min after behavioral examination. They were rapidly decapitated and the brain removed to obtain transverse hippocampal slices (400 μm) from the right dorsal hippocampus. Slices were incubated for 2 hours in oxygenated aCSF (in mM: 119 NaCl, 4.9 KCl, 1.5 MgSO_4_, 2.5 CaCl_2_, 26.2 NaHCO_3_, 1 NaH_2_PO_4_ and 11 Glucose saturated with 95% O_2_, 5% CO_2_), and then were placed in a submerged chamber subfused with aCSF at 35-36°C for recording. A pair of stimulation (bipolar; FHC & Co, ME, USA) and recording electrodes (borosilicate glass pipette filled with aCSF; 5-10 MΩ) was used to evoke and record field excitatory postsynaptic potentials (fEPSP) at the CA1 *stratum radiatum*. Stimulus-response curves were obtained by delivering square pulses (50 μs) at increasing voltages (0-40 V). For synaptic potentiation studies, the test pulse intensity was set at 40% of the maximum fEPSP slope amplitude and sampled once per minute. After a stable baseline response was established, a single 1-s 100-Hz stimulation (high-frequency stimulation, HFS) was delivered to induce long-lasting potentiation. All data sets were normalized to baseline (pre-HFS) values. Slope amplitudes between conditions were compared at three phases of potentiation relative to the baseline: i) post-tetanic potentiation (PTP) measured by averaging during the first 8 min after HFS, early-potentiation measured by averaging during the 10-20 min after HFS, and late-potentiation measured by averaging during 50- 60 min after HFS, measured relative to the 10 min baseline before HFS.

### In vivo Electrophysiology

Male WT (n = 12) and Fmr1-null mice (n = 9) aged 4-6 months were anesthetized by isoflurane inhalation (1.5% with 1 L/min oxygen) and implanted with a microdrive. Either a customized 8-tetrode Open Ephys Flexdrive <u>(http://www.open-ephys.org</u> (Voigts et al., 2013)) or a 4-tetrode customized Versadrive (Neuralynx, Bozeman, MT) was used. Tetrodes were aimed at the dorsal hippocampus (-2.0 mm posterior to bregma, + 1.75 mm lateral from midline). The Flexdrive targeted both hippocampi. The tetrodes were constructed from 18-μm Nichrome or Platinum Iridium wire and the tips were plated with gold or platinum, respectively, to reduce impedances to 200-250 kOhms. The microdrive was secured to the skull using bone screws and dental cement; one screw served as the ground electrode. The mouse recovered for at least a week before electrophysiological procedures began.

Tetrodes were slowly advanced over two weeks until hippocampal single units could be identified. An electrode without single unit activity was selected as a reference. Extracellular action potential signals were amplified 4-12k times and 0.6-6 kHz band-pass filtered. LFP signals from the same electrodes were amplified 1000 times and 0.1-500 Hz band-pass filtered. Electrophysiology data were recorded using a 64-channel Axona recording system (Axona, St. Albans, UK).

### Histology

After data collection concluded, the mice were euthanized by pentobarbital overdose. The mice were transcardially perfused with 4% formalin,brains removed, cryoprotected with 30% sucrose, sliced at 30 μm with a cryostat, and mounted on slides. Cresyl violet stain was used to determine the recording site for each tetrode, which had been targeted to the pyramidal cell layer of CA1.

## Behavioral Procedures

### Constant conditions open-field and disk rotation

Four recording environments were used 1) a small box (28.5 w x 28.5 l x 17 h cm), 2) a large box with black walls (50 w x 50 l x 25 h cm), white acrylic floor, and a checkerboard card attached to one wall, 3) a 10-cm wide, ring with 40-cm outer diameter, and 4) a circular disk-shaped arena (40 cm diameter, 30 cm transparent wall) that could rotate at 0.75 rpm.

Electrophysiology recordings were performed while the mouse explored the small or large boxes, the ring or the disk that was positioned in the same location within the center of the room. A black curtain containing visual orienting cues was placed around the apparatus. The animal’s position was tracked at 30 frames/s using an overhead camera and software (Tracker, Bio-Signal Group Corp., Acton, MA) to detect a pair of infrared diodes attached to the animal’s microdrive. During arena rotation sessions, position was tracked in both the spatial frame of the room and the spatial frame of the arena. Tracking in the arena frame was performed relative to an infrared diode that was attached to the rotating arena but inaccessible to the animals.

### Active place avoidance

Male WT and Fmr1-null mice aged 3-4 months were trained in a hippocampusdependent two-frame active place avoidance task, consisting of a 40-cm diameter arena with a parallel rod floor that rotated at 1 rpm. The position of the animal was tracked as described above. Mice in the trained condition learned the “Room+Arena-“task variant, in which avoiding a 60° sector was reinforced by a constant current foot shock (60 Hz, 500 ms, 0.2 mA) that was scrambled (5-poles) across pairs of the floor rods, triggered by entering the shock zone for more than 500 ms. Additional shocks occurred every 1.5 s until escape. Measures of place avoidance were computed by software (TrackAnalysis, Bio-Signal Group Corp., Acton, MA). The behavioral protocol began with a 10-min session with shock off to habituate the mice to the rotating arena (pretrain). One hour later, three 10-min training sessions (60-min inter-trial interval) followed with the shock turned on. An additional conditioning session “24-h Retest” was performed the following day. Conditions were identical across all sessions except shock was off during pretraining.

## Data Analysis

### Classification of cell types

Manual single unit isolation (Fig. S2) quality was quantified using *IsoI_BG_* and *IsoI_NN_* (Neymotin et al., 2011). Single units were considered sufficiently well isolated for study only if *IsoI_BG_* > 4 bits and *IsoI_NN_* > 4 bits. Single units were classified as complex-spike or theta cells according to published criteria (Fenton et al., 2008; Ranck, 1973) (Fig. S3). Complex-spike cells appear to be pyramidal cells, whereas theta cells are likely local interneurons (Fox and Ranck, 1975). Pyramidal cells were judged to have long-duration waveforms (> 250 μs), low discharge rate (< 2 AP/s) and a tendency to fire in bursts (peak inter-spike interval < 10 ms). Interneurons had short-duration waveforms (< 250 μs), high discharge rate (> 2 AP/s), and were less likely to fire in bursts.

### Characterization of place cells

Cell-specific spatial firing rate maps were created by calculating the total number of spikes observed in each 1.25 × 1.25 cm location, divided by the total time the mouse was in the location. Three qualities of spatial firing were computed from the firing rate map: the overall firing rate is the total spikes a cell discharged divided by the total recording time; spatial coherence describes the local smoothness of the firing rate distribution (Muller and Kubie, 1989); and spatial information content describes the reduction in uncertainty of the mouse’s position in the firing rate map given a particular firing rate (Skaggs et al., 1993). Single units were classified as place cells if the overall discharge rate was (0.1-2 AP/s), the spatial coherence (z score) was > 0.4, and the information content was > 0.4 bits/spike. The spatial similarity of two firing rate maps was computed as Fischer’s z-transformation of Pearson’s correlation for the firing rates in corresponding pixels of the two maps. The proportion of pixels in which the cell discharged was also computed.

### Overdispersion

Overdispersion is the variance of a cell’s standardized firing rates for passes through the firing field (Fenton et al., 2010). The standardized firing rate (*z*) was computed for each 5-s interval as:

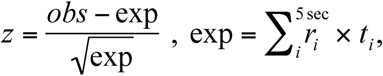
where *obs* is the number of observed action potentials and *exp* is the expected number on the assumption of Poisson firing. The expectation is the sum of the product of the time spent (*t_i_*) in a location during time interval *i* and the time-averaged rate at that location (*r_i_*) computed for all locations that were visited during the 5 seconds. Note that *exp* = 0 and *z* is undefined if the mouse does not visit the firing field during a 5-s interval. Only 5-s intervals when the mouse sampled a cell’s firing field well were studied. These were identified by *exp* ≥ average rate for the cell (Fenton et al., 2010).

### Functional coupling of cell pairs

The functional coupling of spike trains from pairs of cells was estimated using Kendall’s correlation (Neymotin et al., 2017). The time series was generated by counting the number of spikes the cell fired during each time interval. We examined gamma (25 and 40 ms), theta (125 and 250 ms), as well as 1-s and 5-s (time to cross place field) intervals. Distributions of the population of pair-wise correlations within an ensemble of cells are called PCorr, which is the vector of pairwise correlations. PCorr captures higher-order network correlations, as weak pair-wise correlations imply strong network states (Schneidman et al., 2006, Neymotin et al., 2017). We used PCo, the Pearson correlation of a pair of PCorr vectors to describe the recurrence of these network states (Neymotin et al., 2017).

### Frame-specific spatial firing analysis

During arena rotation, spatial discharge of a neuron can be represented in either the stationary spatial frame of the room or the rotating spatial frame of the arena. At each moment, discharge can signal location in one frame or the other. To decode which spatial frame is being represented by location-specific CA1 discharge, we computed the momentary positional information *I_pos_* as described previously (Kelemen and Fenton, 2010). *I_pos_*(*t*) estimates the location-specific firing of a cell during a brief time interval (Δt = 133 ms). It is defined as:
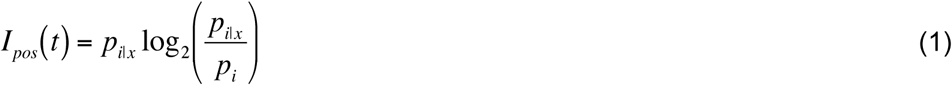

*p_i_* is the probability of the cell firing *i* spikes during the interval; *p_i|x_* is the probability of firing *i* spikes if the mouse is in location *x*. Although *I_pos_*(*t*) can be positive or negative, the absolute value is large whenever the number of spikes observed at the location is distinct or “surprising” compared to the location-independent probability of observing the same number of spikes. The value 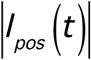 is given the shorthand, *I_pos_*.

A cell’s spatial frame preference was estimated by first calculating *I_pos_* separately for each spatial frame, then by computing the difference Δ*I_pos_* = *I_pos_*_(*room*)_− *I_pos_*_(*arena*)_.

To estimate ensemble Δ*I_pos_*(*t*), we first computed the frame-specific sum of the *I_pos_*(*t*) values at each moment. The ensemble difference Δ*I_pos_* between the room and arena frame was computed at each time interval to estimate the momentary frame preference in the ensemble discharge. The Pearson correlation between the time series of ensemble Δ*I_pos_* values and the time series of an individual cell’s Δ*I_pos_* values was used to estimate the coordination between an ensemble and a single cell’s frame-specific fluctuations in positional information. This coordination was estimated for each cell in the ensemble by computing the ensemble Δ*I_pos_* time series after leaving the cell out of the ensemble.

### Spike-field coordination

Phase-frequency discharge probability plots were computed to characterize the likelihood that a cell’s discharge is phase-organized by frequency-specific oscillations in the LFP. The LFP signal was convolved with a group of complex Morlet wavelets in the logarithmic range between 2 and 100 Hz. The instantaneous phase of the frequency band-specific LFP signals was obtained from the complex time series. The oscillation phase discharge probability was computed independently for each frequency band. The frequency-specific phase distribution of spiking was normalized by dividing the discharge distribution by the total number of action potentials in a given recording.

## Statistical Analysis

Average ± s.e.m. values are reported throughout the manuscript. Statistical evaluation of data in figures is presented in the legend. Parametric comparisons of the two genotypes were performed by t test; comparisons of binomial proportions used the proportions z test. Multiple factor comparisons were performed by ANOVA, with repeated measures, followed by Tukey post-hoc tests as appropriate. Statistical comparisons between distributions of correlations were performed on Fischer z-transformed correlation values. Significance was set at < 0.05.

## Supplemental Information

Supplemental Information includes five figures and can be found with this article online.

## Acknowledgments

Supported by Simons Foundation grant SFARI 294388 and NIH grant R01MH099128 to A.A.F. and R21NS091830 to J.M.A, NIMH studentship MH96331-5 to Z.N.T., and a Canadian Institutes of Health Research fellowship to F.T.S.

## Author Contributions

FTS and ZNT collected and analyzed place cell data, ZNT, DD, AAF analyzed spike train data, DD and AAF created software and hardware for data collection, JMA, BMC, AAF collected and analysed *ex vivo* data, AAF supervised research and wrote the manuscript with contributions from all authors.

